# Distributed correlates of visually-guided behavior across the mouse brain

**DOI:** 10.1101/474437

**Authors:** Nicholas A. Steinmetz, Peter Zatka-Haas, Matteo Carandini, Kenneth D. Harris

## Abstract

Behavior arises from neuronal activity, but it is not known how the active neurons are distributed across brain regions and how their activity unfolds in time. Here, we used high-density Neuropixels probes to record from ~30,000 neurons in mice performing a visual contrast discrimination task. The task activated 60% of the neurons, involving nearly all 42 recorded brain regions, well beyond the regions activated by passive visual stimulation. However, neurons selective for choice (left vs. right) were rare, and found mostly in midbrain, striatum, and frontal cortex. Those in midbrain were typically activated prior to contralateral choices and suppressed prior to ipsilateral choices, consistent with a competitive midbrain circuit for adjudicating the subject’s choice. A brain-wide state shift distinguished trials in which visual stimuli led to movement. These results reveal concurrent representations of movement and choice in neurons widely distributed across the brain.

---

Many studies have examined the role of individual brain regions in sensory processing and decision making^1–5^, but it remains unclear whether these processes are implemented by specialized regions or by distributed circuits. For example, activity in sensory cortices correlates not only with sensory stimuli but also with motor planning^6,7^, movement^8^, upcoming behavioral reports^9–14^, spatial attention^15,16^ and reward^17^. Similarly, in motor regions such as the deep layers of the superior colliculus^18^, activity correlates with aspects of decision making and other cognitive functions^5,19–23^. The brain regions implicated to date in sensory-guided behavior include neocortex, basal ganglia, thalamus, and midbrain. These regions are widely and reciprocally connected^24–26^, suggesting that any particular aspect of sensory, decision, or motor processing that involves neurons in one region may also involve the coordinated activity of neurons across others. This hypothesis is consistent with evidence that task engagement^27^, rewards^28^, and spontaneous beahviors^29^ modulate activity across wide cortical and subcortical territories. However, testing this hypothesis would require a systematic survey of how neurons across the brain correlate with goal-directed behavior. Such a survey has now become possible thanks to the introduction of high-density probes^30^ able to record from hundreds of neurons simultaneously across multiple regions of mouse brain.

## Widespread activity accompanies visually guided decisions

We trained mice in a head-fixed two-alternative unforced choice (2AUC) task^31^, in which they indicated whether a stimulus on the left or right had higher contrast by turning a wheel with their forepaws (Fig 1a,b), or indicated the lack of either stimulus by holding the wheel still. Trained mice successfully chose Left or Right according to contrast difference, and withheld movements when no stimuli were present (Fig 1c). They performed the task accurately for high-contrast single stimuli (i.e. when the other stimulus was absent; 88 ± 9% correct, mean ± s.d., n = 39 sessions, 10 mice), but less accurately in more challenging conditions: with low-contrast single stimuli (65 ± 20% correct mean±s.d); or with competing stimuli of similar contrast (66 ± 11% correct mean±s.d., on trials with high vs. medium or medium vs. low contrast). Reaction times were slower for trials with two competing stimuli than with single stimuli of the same contrast (Fig 1d, p<10^-4^, multi-way ANOVA), suggesting that mice selected a choice by resolving a competition between sensory information from the two stimulus locations.

**Figure 1.**
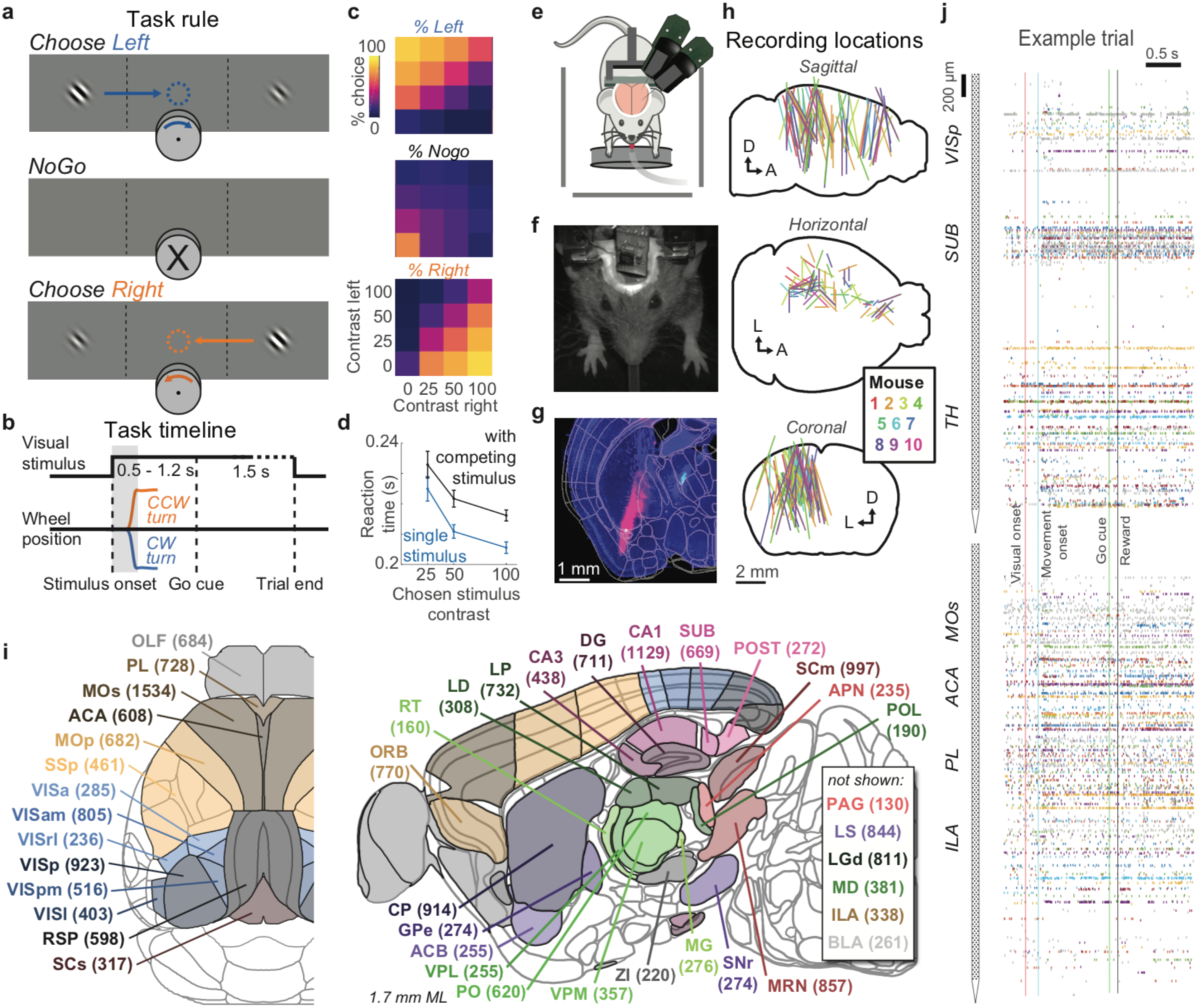
Recordings in 42 brain regions during a two-alternative unforced choice task. **a,** Mice earned water rewards by turning a wheel to indicate which of two visual gratings had higher contrast, or not turning if no stimulus was presented. When stimuli had equal contrast, a Left or Right choice was rewarded with 50% probability. Grey rectangles with dashed dividers indicate the three computer screens surrounding the mouse. Arrows (not visible to the mouse) indicate the coupled movement of the visual stimulus with the wheel, and the colored dashed circle (not visible to the mouse) indicates the stimulus location at which a reward was delivered. **b,** Timeline of the task. Subjects were free to move as soon as the stimulus appeared, but the stimulus was fixed in place and rewards were unavailable until after an auditory go cue. If no movement was made for 1.5 s after the go cue, a NoGo was registered. The grey region is the analysis window, from 0 to 0.4 s after stimulus onset. **c,** Average task performance across subjects, n=10 subjects, 39 sessions, 9,538 trials. Colormaps depict the probability of each choice given the combination of contrasts presented. **d,** Reaction time as a function of stimulus contrast and presence of competing stimuli. **e,** Mice were head-fixed with forepaws on the wheel while multiple Neuropixels probes were inserted for each recording. **f,** Frontal view of subject performing the behavioral task during Neuropixels recording, with forepaws on wheel and lick spout for acquiring rewards. **g,** Example electrode track histology with atlas alignment overlaid. **h,** Recording track locations as registered to the Allen Common Coordinate Framework 3D space. Each colored line represents the span recorded by a single probe on a single session, colored by mouse identity. D, dorsal; A, anterior; L, left. **i,** Summary of recording locations. Recordings were made from each of the 42 brain regions colored on the top-down view of cortex (left) and sagittal section (right). For each region, number in parentheses indicates total recorded neurons. **j,** Spiking raster from an example individual trial, in which populations of neurons were simultaneously recorded across visual and frontal cortical areas, hippocampus, and thalamus. For abbreviations, see Extended Data Table 1.

We used Neuropixels probes^30,32^ to record from ~30,000 neurons in 42 brain regions during task performance, and found that nearly 60% of them were modulated during the task (Fig 1e-j). Inserting two or three probes at a time yielded simultaneous recordings from hundreds of neurons in multiple brain areas during each recording session, for multiple days per mouse (n=92 probe insertions over 39 sessions in 10 mice, Fig 1h-j). We identified the firing times of individual neurons using Kilosort^33^ and determined their anatomical locations by combining electrophysiological features with histological reconstruction of fluorescently-labeled probe tracks (Fig 1g, see Methods). Across all sessions we recorded from 29,134 neurons (n=747 ± 38 neurons per session, mean±s.e.), of which 22,458 were localizable to one of 42 brain regions. Of these, 60.0% (13,467) neurons distributed in all regions had detectable modulation of firing rate during the time between stimulus onset and movement and were included in subsequent analyses. The dataset collected for this study will be shared publicly at the time of publication.

Trial onset was followed by increased neuronal activity in nearly all recorded regions (Figure 2a-c). Neurons were diverse in the times they became active during trials, with timing differences both within and between brain areas. Activity emerged first in visual regions contralateral to presented stimuli, and soon afterwards spread to most recorded regions, including areas ipsilateral to the stimulus (Figure 2f-g). This widespread activity began prior to the earliest detectable movement onset in most regions (Fig 2d). Similarly widespread activity was observed following reward delivery (Fig 2e).

**Figure 2.**
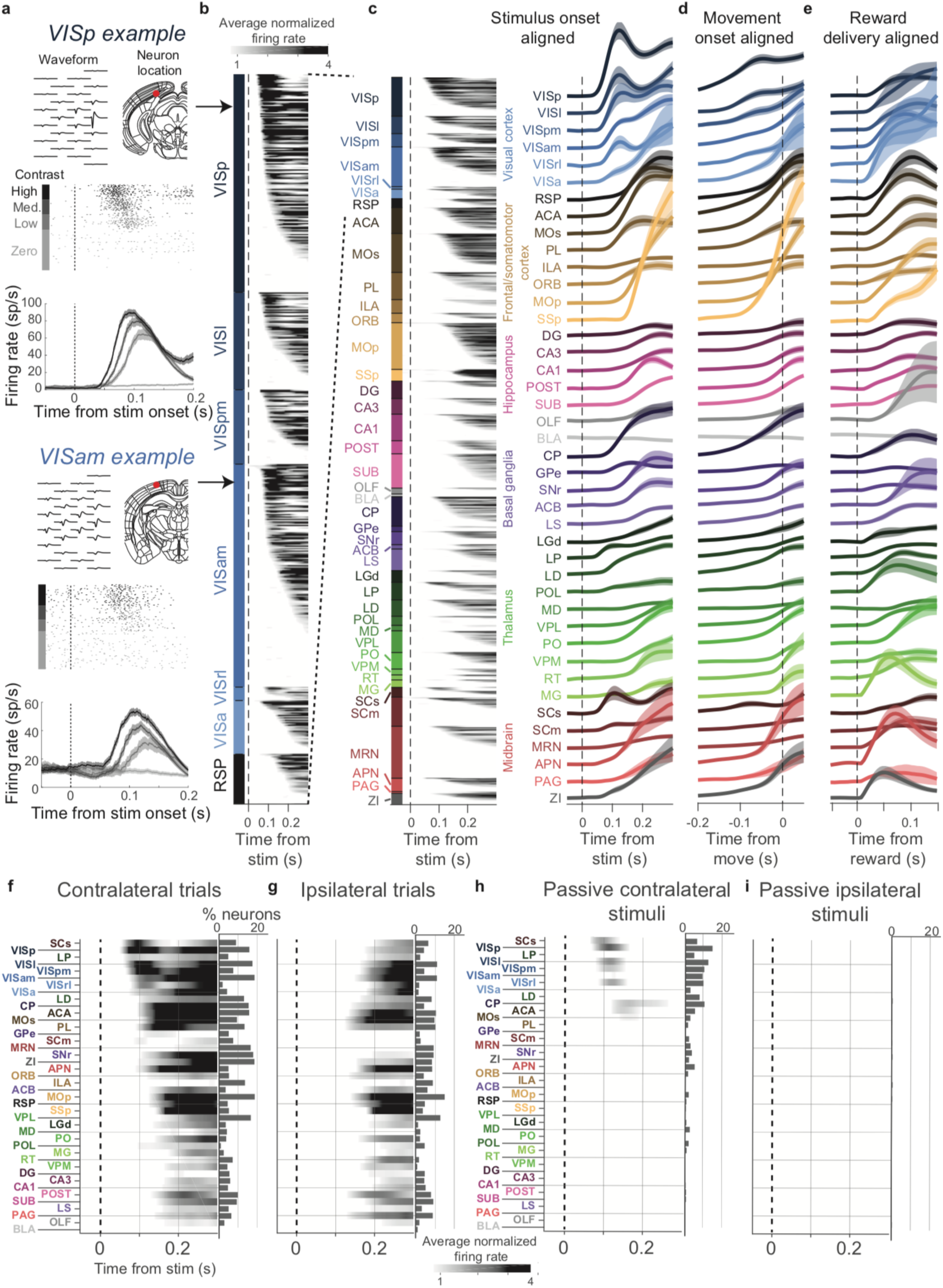
Propagation of activity during task performance. **a,** Activity of example neurons in VISp and VISam, showing the neuron’s waveform and anatomical location (*top*), rasters sorted by contralateral contrast (*middle*), and peri-stimulus time histograms (PSTHs, smoothed with 30 ms causal half-Gaussian) for each of the four contralateral contrasts (*bottom*). PSTH shaded regions: ± s.e. across trials. **b,** Colormap showing PSTHs of all highly-activated neurons (p<10^-4^ compared to pre-trial activity) in posterior cortex, vertically sorted by firing latency. Gray scale represents average normalized firing rate across all trials with contralateral stimuli and with movement. **c,** Left: colormap showing PSTHs of neurons from all recorded brain regions (left), and curves showing mean stimulus-aligned PSTH of all neurons in each area (right). Shaded regions: ± s.e. across neurons. **d,** Curves showing mean PSTH of the same neurons on the same trials, aligned to movement onset. **e,** Curves showing mean PSTH of the same neurons, aligned reward delivery following successful NoGo trials (i.e. trials with no visual stimulus and no overt movement). **f,** Left: colormap showing average PSTH for all neurons in each region on trials with contralateral stimuli and contralateral choices. Right: percentage of neurons in each area significantly active on these trials. **g,** Same as f, for trials with ipsilateral stimuli and ipsilateral choices. **h,** Same as f, for passive presentation of contralateral visual stimuli, excluding any trials with wheel movements. **i,** Same as h, for ipsilateral visual stimuli.

The widespread activity following trial onset was not a direct consequence of the visual stimulus (Fig 2h,i). To isolate circuits responding to the visual stimulus itself, we followed the behavioral sessions with passive replay periods, recording the same neurons and presenting the stimuli the subjects had just experienced in the task, but without the opportunity to earn rewards. In contrast to the widespread activity evoked in the task, responses to these passive visual stimuli were limited to contralateral visual cortex, visual thalamus, striatum and superficial superior colliculus, along with sparse neurons in parts of in frontal cortex, midbrain, and thalamus (Fig 2h). In the passive condition, moreover, we observed no activity in the hemisphere ipsilateral to the stimulus (Fig 2i). Activity across the brain therefore differed greatly depending on whether mice were engaged in the task: although the visual stimulus itself generated activity in a restricted set of regions, this response spread widely across the brain when the stimulus triggered a movement.

## Engagement in the task is accompanied by a brain-wide state change

A possible clue to how task engagement gates the spread of activity comes from its effect on pre-stimulus activity (Figure 3a,b). We considered activity in the 0.2 s prior to stimulus onset, when the subjects waited motionless for the next trial to begin. In this pre-stimulus interval, the firing rate of many regions differed markedly between passive and task conditions (Figure 3b). Relative to the passive condition, pre-stimulus firing rates measured during the task were lower in visual cortex and visual thalamus, but higher in several subcortical regions, including basal ganglia and midbrain structures (p<0.05 with Bonferroni correction, nested ANOVA).

**Figure 3.**
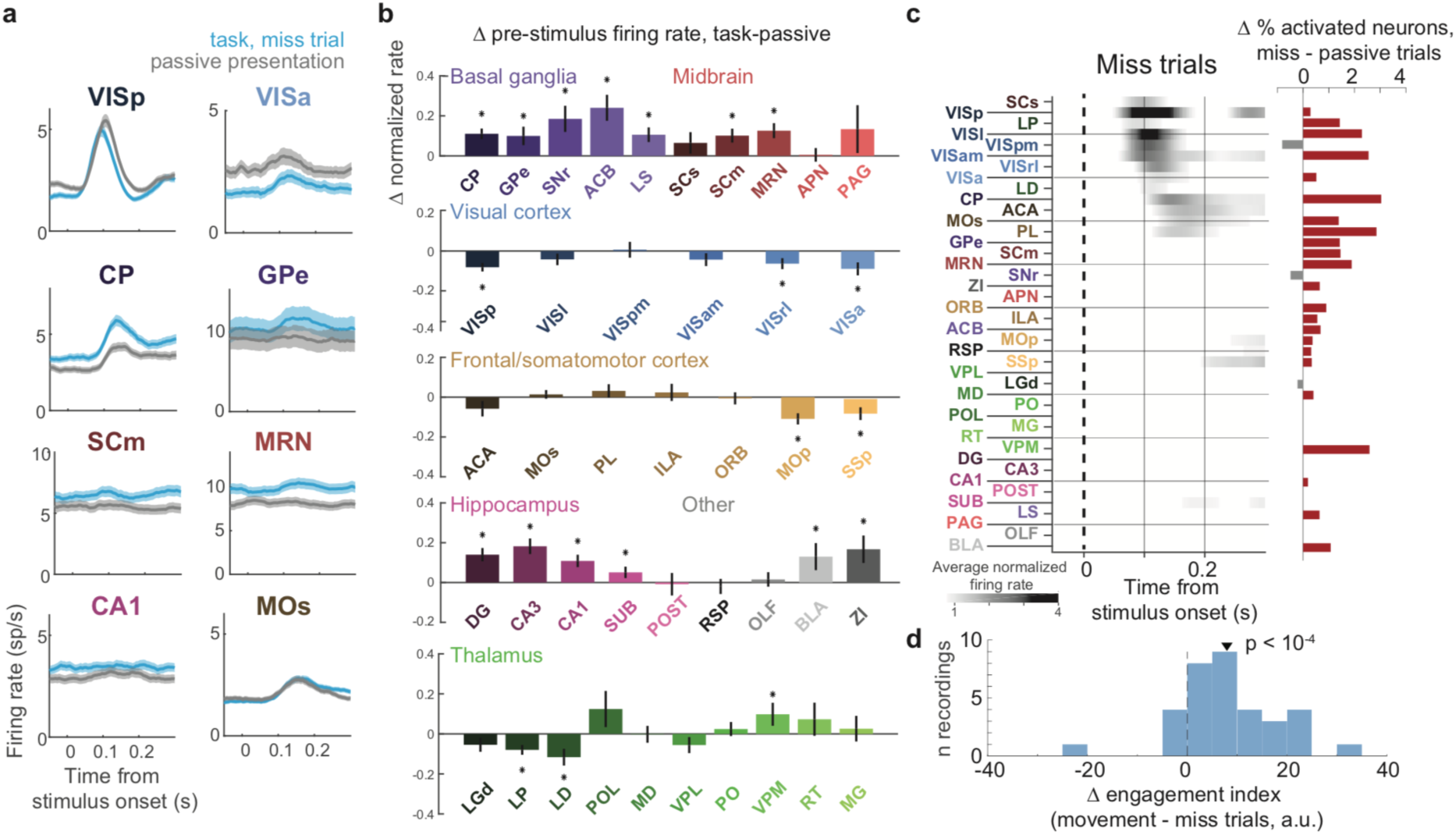
Contextual modulation of pre-stimulus firing rates between passive and active task conditions. **a,** Comparison of population average spiking activity for several brain regions, for trials when contralateral stimuli were presented but subjects did not make a choice (i.e. ‘miss’ trials, blue) and for stimulus presentations during the passive condition (grey). Visual stimulus contrasts were matched between the two conditions so that differences in activity do not reflect differing visual drive. **b,** Difference in log-transformed pre-stimulus firing rate between neurons recorded during task performance and the same neurons recorded during the passive condition. Positive numbers indicate higher firing rate in pre-stimulus periods during the task than in passive. *, p<0.0012. **c,** Left, As in Fig 2f, but for miss trials. Right, the excess proportion of neurons active in this condition relative to the passive condition, for matched contrast stimuli. **d,** Histogram of differences in pre-stimulus engagement index for movement versus miss trials for each recording. Inverted triangle represents the mean value across recordings (mean = 8.42 a.u.).

Activity on miss trials within the task constituted a state intermediate between engaged and passive (Fig 3c,d). Following stimulus onset on miss trials (stimulus presented but no movement), activity spread to more contralateral regions than in the passive context, and a greater number of neurons became active (p<10^-7^, Fisher exact test, matched stimulus contrasts; Fig 3c), although activity on miss trials was not as widespread as in trials with movement. Moreover, pre-stimulus activity resembled passive state activity more on miss trials than on movement trials. For each trial we computed an “engagement index” by projecting the *N_neurons_*-dimensional vector of pre-stimulus firing rates from each trial onto the vector of average pre-stimulus differences between all active and all passive trials. The engagement index was significantly greater for movement trials than miss trials (t-test, p<10^-4^; Fig 3d; matched stimulus contrasts), indicating that brain-wide state measured prior to stimulus onset exhibited a continuum, with the state prior to miss trials intermediate between passive and movement conditions. We hypothesize that this state change reflects an enhanced excitability of certain structures (such as basal ganglia and midbrain, which showed stronger activity), that amplifies stimulus-driven activity during the task.

## Neurons encoding choices are rare but distributed

While most neurons active prior to the movement fired equally for left and right choices, rare neurons showed strong correlates of choice direction (Figure 4a). To distinguish neurons genuinely encoding a choice from neurons encoding stimulus contrast, we employed a statistical method that disentangled these two correlated quantities, exploiting the fact that difficult stimuli elicited variable choices, with variable timings (cf. Figures 1c,d). We first fitted each neuron's activity with a sum of kernel functions time-locked to stimulus presentation and to movement onset^34^. We fit six stimulus-locked kernels - one for each of three possible contrast values on each side - which captured variations in amplitude and timing of the visual activity driven by different stimuli (cf. Figure 2a). We fit two movement-locked kernels: one triggered by a movement in either direction (‘Move’), and one capturing differences in activity between Left and Right movement directions (‘Choice’). To avoid overfitting, we devised a kernel estimation method based on reduced-rank regression, leveraging the large number of recorded neurons to identify a low-dimensional representation that accurately fit the activity of neurons across brain regions and recordings.

**Figure 4.**
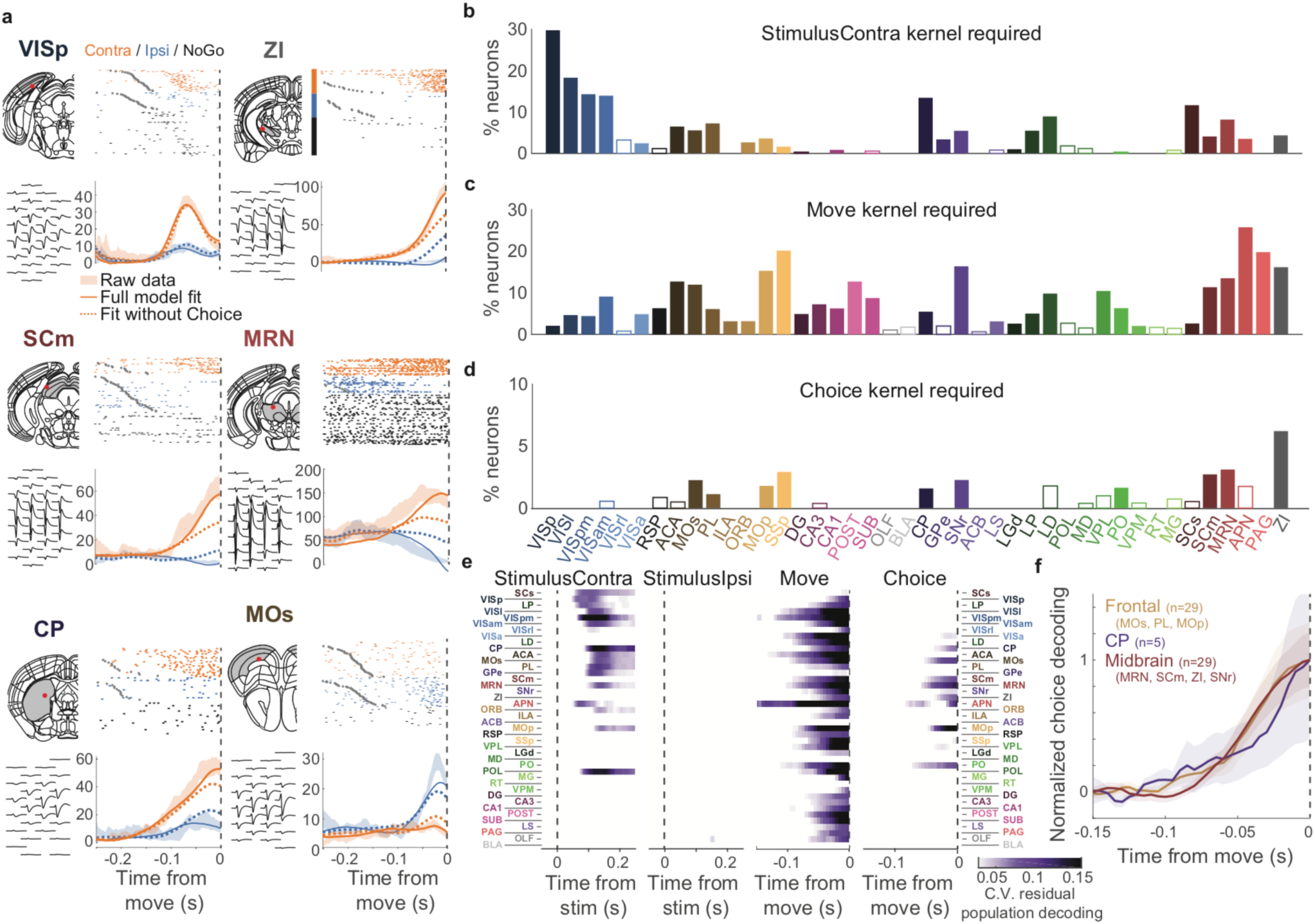
Regression analysis reveals widespread representation of movement occurrence but rare representation of choice. **a,** Example neurons from indicated brain regions, with registered atlas location (upper left) and waveform (lower left). Rasters (upper right) show each neuron’s activity aligned to movement onset, with trials color coded and vertically arranged by the subject’s choice (grey circles represent stimulus onsets; for NoGo trials, spikes are aligned to the expected movement time for the contrasts shown). Lower right panels show PSTHs (shaded regions indicating mean ± s.e.), cross-validated predictions of the full kernel model (solid lines, derived from stimulus identity and timing, choice identity and timing), and model missing the Choice kernel (dashed lines). **b,** Proportion of individual neurons for which accurate prediction required the Contralateral Stimulus kernel (cross-validated variance explained > 2%). Empty bars indicate those for which the number of observed neurons was < 5. **c,** As in b, but for the Move kernel. **d,** As in b, but for the Choice kernel. Note the difference in y-scale from b and c. **e,** Population decoding of Stimulus Contra identity from residual activity in each area after fitting a model including all other kernels. Subsequent three panels depict the same analysis for the other three kernel types. **f,** Time course

To determine whether a neuron encoded stimulus, movement, or choice, we asked whether the corresponding kernel was necessary to predict its activity. We first fit a model including all kernels except the one to be tested, and asked whether adding the test kernel improved this fit for held out data. For example, a representative neuron in primary visual cortex (VISp) showed strong firing rate differences between Left and Right choices (Figure 4a, first panel, shaded regions), but these were time-locked to the visual stimuli and could be accurately predicted with or without the choice kernel (solid and dashed lines), suggesting the neuron only encoded visual contrast. However, other neurons which increased their firing time-locked to movements of a particular direction could not be predicted without the Choice kernel (Figure 4a, other panels). Consistent with the previous comparison of trial and passive stimulus responses (Fig. 2), we found encoding of upcoming movement (independent of direction) in nearly all brain regions (Figure 4c), but encoding of the contralateral stimulus primarily in visual and frontal cortex, visual thalamus, striatum, and superficial superior colliculus (Figure 4b).

Neurons encoding the upcoming choice were rare, and limited to a distinctive set of regions in cortex, striatum, and midbrain (Fig 4d). Neurons for which the Choice kernel was required to accurately predict activity were found in frontal cortex (MOs, PL, and MOp), striatum (CP), and midbrain (SNr, SCm, MRN, and ZI). Although many neurons in these regions had strong choice correlates (e.g. Fig. 4a), choice-selective neurons accounted for a small fraction of cells in these regions (at most 6% in ZI, notice different scale between Fig. 4d and Fig. 4b,c). This small population, however, sufficed to predict the mouse’s impending choice prior to movement onset (Fig 4e; last panel). The time-course of choice decoding from population activity was not significantly different in these regions (two-way ANOVA, p > 0.05; Fig 4f).

## Choices are encoded differently in midbrain and forebrain

Although the encoding of choice emerged simultaneously in midbrain and forebrain, it was encoded differently in these regions (Figure 5). Nearly all (53/54, 98%) choice-selective neurons in midbrain (MRN, SCm, SNr, and ZI) preferred contralateral choices (Fig. 5a, top, and Fig. 5b,c). By contrast, neurons in forebrain (MOs, PL, MOp, and CP) could prefer either choice (29/48, 60% preferred contralateral; Fig. 5a, bottom, and Fig. 5e,f). This difference between midbrain and forebrain was significant (p=<10^-5^, Fisher’s exact test; Figure 5e,f). Midbrain neurons, moreover, exhibited directionally-opposed activity: their activity increased before one choice and decreased below baseline before the other (29/54, 54%; Figure 5d), as though competitively inhibited prior to ipsilateral choices by other neurons preferring that choice. Forebrain neurons typically exhibited a qualitatively distinct encoding: neurons in one hemisphere became active prior to both left and right choices (10/48, 21% suppressed for non-preferred choice). This difference between midbrain and forebrain was also significant (p<10^-3^, Fisher’s exact test; Figure 5f).

**Figure 5.**
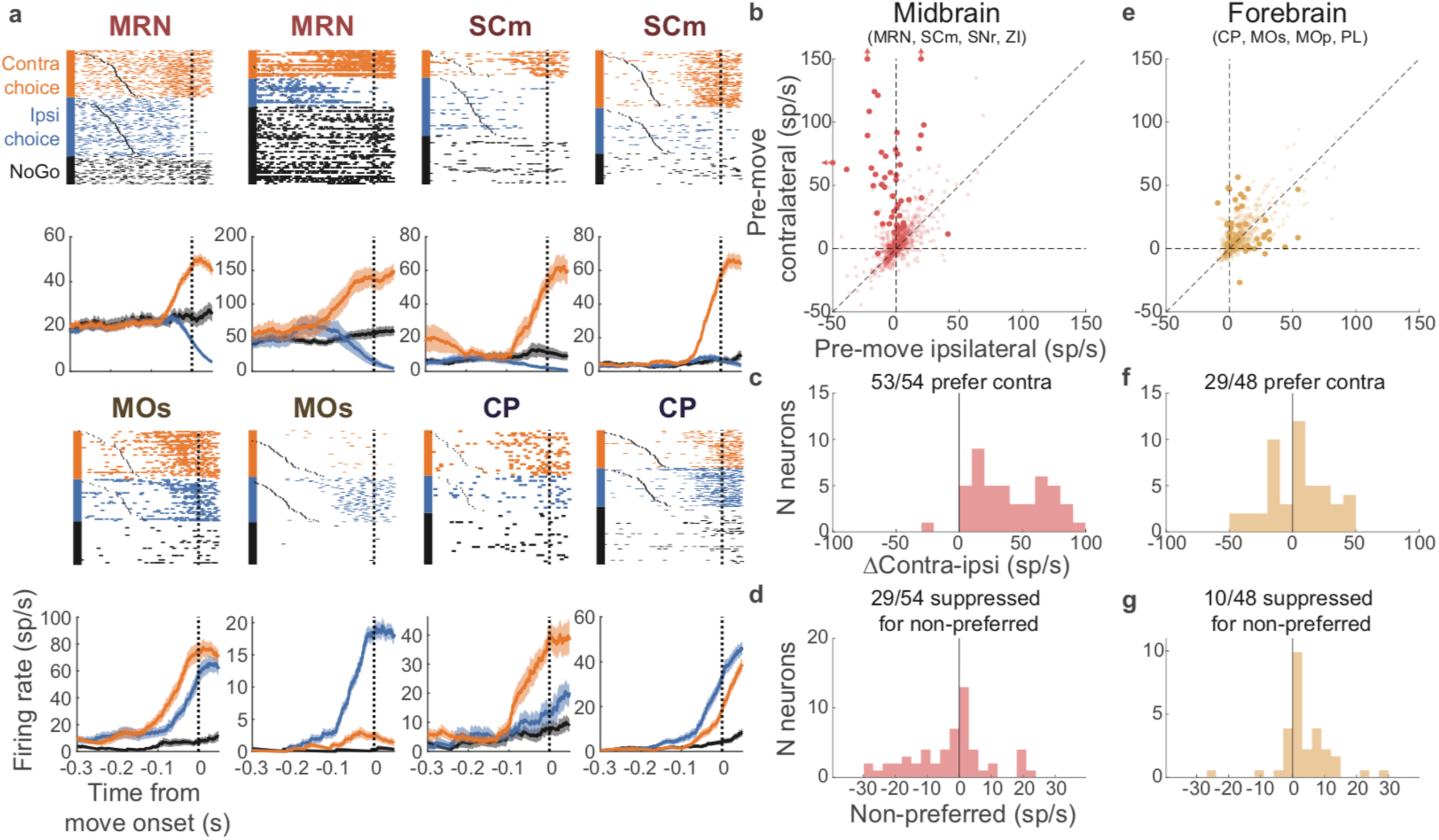
Choices are encoded distinctly in midbrain and forebrain neurons. **a,** Example neurons recorded in the midbrain (top row) and forebrain (bottom row). Conventions as in Fig 4a. **b,** Scatter plot of activity of individual midbrain neurons at movement onset vs baseline activity, for trials with contralateral versus ipsilateral choices (estimated from the kernel model). Darker points represent neurons with significant choice encoding (c.f. Fig 4d). **c,** The difference in response amplitude between contralateral and ipsilateral choice trials, for midbrain neurons with significant choice encoding. **d,** Amplitude on non-preferred trials relative to baseline. **e, f, g,** as in b, c, d but for forebrain neurons.

## Discussion

We found that neurons across many disparate brain regions were activated when mice performed the task. Nevertheless, the key signals required for task performance were encoded by smaller networks. Visual onset was encoded by neurons in superficial SC, visual cortex, visual thalamus, frontal cortex, striatum, and sparse neurons elsewhere. An even smaller subset of neurons in frontal cortex, basal ganglia, and midbrain structures encoded left/right choices. Choice-selective neurons in the midbrain but not forebrain exhibited competitive coding, with contralateral choices preceded by increases in rate, and ipsilateral choices preceded by decreases. Taken together, these observations indicate that signals reflecting an impending choice emerge across a coordinated network of neurons in midbrain, frontal cortex, striatum and thalamus; and that competitive circuits within the midbrain may ultimately adjudicate the decision.

In contrast, we saw a global but nonspecific brain activation around the time of movement onset. This activation may largely reflect a corollary discharge, downstream of the decision to initiate a movement^35,36^. Although observational results cannot alone determine causal involvement of a particular brain region in the task, another study^37^ using the same task confirms this hypothesis at least for some parts of dorsal cortex: while optogenetic inactivation of visual and frontal cortex affected the mouse’s performance, inactivation of primary somatomotor cortex and retrosplenial cortex did not despite the robust pre-movement activity we observed in these regions.

The hypothesis that a network of midbrain areas adjudicates competition between two alternative choices is consistent with work describing roles for SC in decision-making in rats^5^ and primates^22,38^. The competitive interaction could arise from mutually inhibitory circuitry within the SC^39^, or from intra-midbrain inhibitory projections from SNr to SC^4,40^. Although the MRN and ZI have been less well-studied than SC, recordings in primates during arm movements suggest that MRN and SC carry similar signals^41^. Our results suggest that these midbrain nuclei are involved in action selection, not just motor execution. Their near-exclusive firing for contralateral choices – executed through both forelimbs – suggests a high-level representation of the choice, rather than a low-level representation of muscle contractions. Nevertheless, our results argue against SC and MRN as the sole locus for the decision at the end of a feedforward pathway. Choice-predictive signals emerged in frontal cortex and striatum at approximately the same time, albeit with a primarily non-competitive encoding. These data are therefore compatible with a view that midbrain neurons mediate competition between wider recurrently-connected circuits in each hemisphere.

The idea that actions are selected through recurrent self-exciting subnetworks that are coupled by inter-network inhibition has been studied in computational models^42,43^. In these models, recurrent excitation causes threshold behavior: a self-exciting subnetwork receiving inputs proportional to the evidence in favor of a choice generates all-or-none responses when this evidence exceeds a threshold; and mutual inhibition between two such subnetworks allows the choice with more evidence to suppress the choice with less. Our data are consistent with a choice mechanism consisting of a recurrent circuit in each hemisphere, spanning frontal cortex, basal ganglia, and midbrain nuclei, with a net self-excitation in each side, and coupled by inhibition within the midbrain (Fig 6a). Engagement in the task correlated with a boost in pre-stimulus activity in these same structures, suggesting that engagement is accompanied by tonic input specifically to these networks. This input could come from dopamine or other neuromodulator systems^44^, and might reflect either a task-specific command, or a general increase in arousal. We hypothesize that only with this tonic input are sensory stimuli sufficient to bring the network past threshold, explaining why movements follow sensory stimuli only when the subjects are engaged in the task (Fig 6b-d).

**Figure 6.**
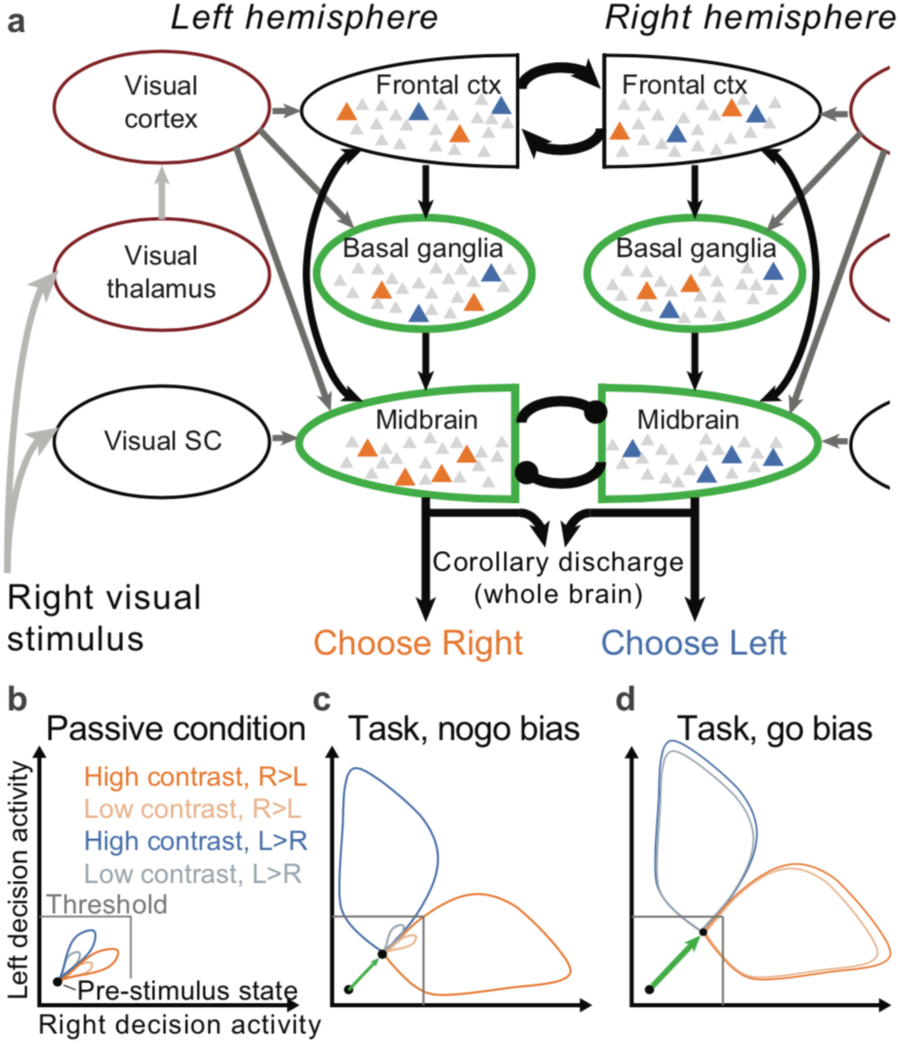
Hypothetical circuit organization and dynamics. **a,** Visual activity first arises in in SCs, visual thalamic nuclei and visual cortex, then propagates to a self-exciting decision network containing frontal cortex, basal ganglia, and midbrain circuits. Competition between left and right choices is adjudicated by inhibitory interactions within the midbrain. Arrowheads represent excitatory connections, and circles represent inhibition. Arrow color represents time of activity during the task. Green outlines represent areas with boosted pre-stimulus activity during the task; red those with suppressed. **b-d**, A cartoon dynamical model of decision initiation. X- and y-axes represent activity in right and left decision networks. The three panels represent dynamical trajectories in the passive condition, task condition on NoGo-biased trials, and task condition on movement-biased trials, characterized by progressively higher baseline activity in both decision networks. Stimulus onset increases activity in the two decision networks according to its contrast in the corresponding side. In the passive condition (b), baseline activity is so low that activity cannot cross threshold even for high-contrast stimuli. In the NoGo-biased task condition (c), high-contrast stimuli are sufficient to bring one decision network to threshold, which then suppresses activity in the other, but low contrast stimuli do not cause either network to cross threshold. In the movement-biased task condition (d), even low-contrast stimuli are sufficient to bring one or other decision network to threshold.

In conclusion, the correlates of choice in a single task are distributed and near-simultaneous across disparate brain systems. This finding motivates a change in focus in the search for the neural basis of behavior: from specialized brain regions to coordinated circuits of individual neurons distributed across the brain.

## Acknowledgements

We thank M. Pachitariu and P. Shamash for analysis tools; H Forrest for help with data preprocessing; C Reddy, M Wells, L Funnell for help with mouse husbandry, training, and histology; R Raghupathy for help with histology; C.P. Burgess for help with experimental apparatus. We thank T. Harris and B. Karsh for support with Neuropixels recordings. We thank M. Häusser, A.J. Peters and S. Schroeder for feedback on the manuscript. This project was funded by the European Union’s Marie Skłodowska-Curie program (656528), the Human Frontier Sciences Program (LT001071/2015-L), the Wellcome Trust (205093, 102264), the European Research Council (694401), the Gatsby Foundation (GAT3531), and the Simons Foundation (325512). M.C. holds the GlaxoSmithKline/Fight for Sight Chair in Visual Neuroscience.

## Author contributions

N.A.S., M.C., and K.D.H. conceived and designed the study. N.A.S. collected data. N.A.S. and P.Z-H. analyzed data. N.A.S., M.C., and wrote the manuscript.

## Author information

Correspondence and requests for materials should be addressed to.

## Methods

Experimental procedures were conducted according to the UK Animals Scientific Procedures Act (1986) and under personal and project licenses released by the Home Office following appropriate ethics review.

### Subjects

Experiments were performed on male and female mice, between 11 and 46 weeks of age (Extended Data Table 2). Multiple genotypes were employed, including: Ai95;Vglut1-Cre (B6J.Cg-Gt(ROSA)26Sor^tm95.1(CAG-GCaMP6f)Hze^/MwarJ crossed with B6;129S-Slc17a7^tm1.1(cre)Hze^/J), TetO-G6s;Camk2a-tTa (B6;DBA-Tg(tetO-GCaMP6s)2Niell/J crossed with B6.Cg-Tg(Camk2a-tTA)1Mmay/DboJ), Snap25-G6s (B6.Cg-Snap25^tm3.1Hze^/J), Vglut1-Cre, and wild-type (C57Bl6/J). None of these lines are known to exhibit aberrant epileptiform activity^45^.

**Extended Data Table 1.**
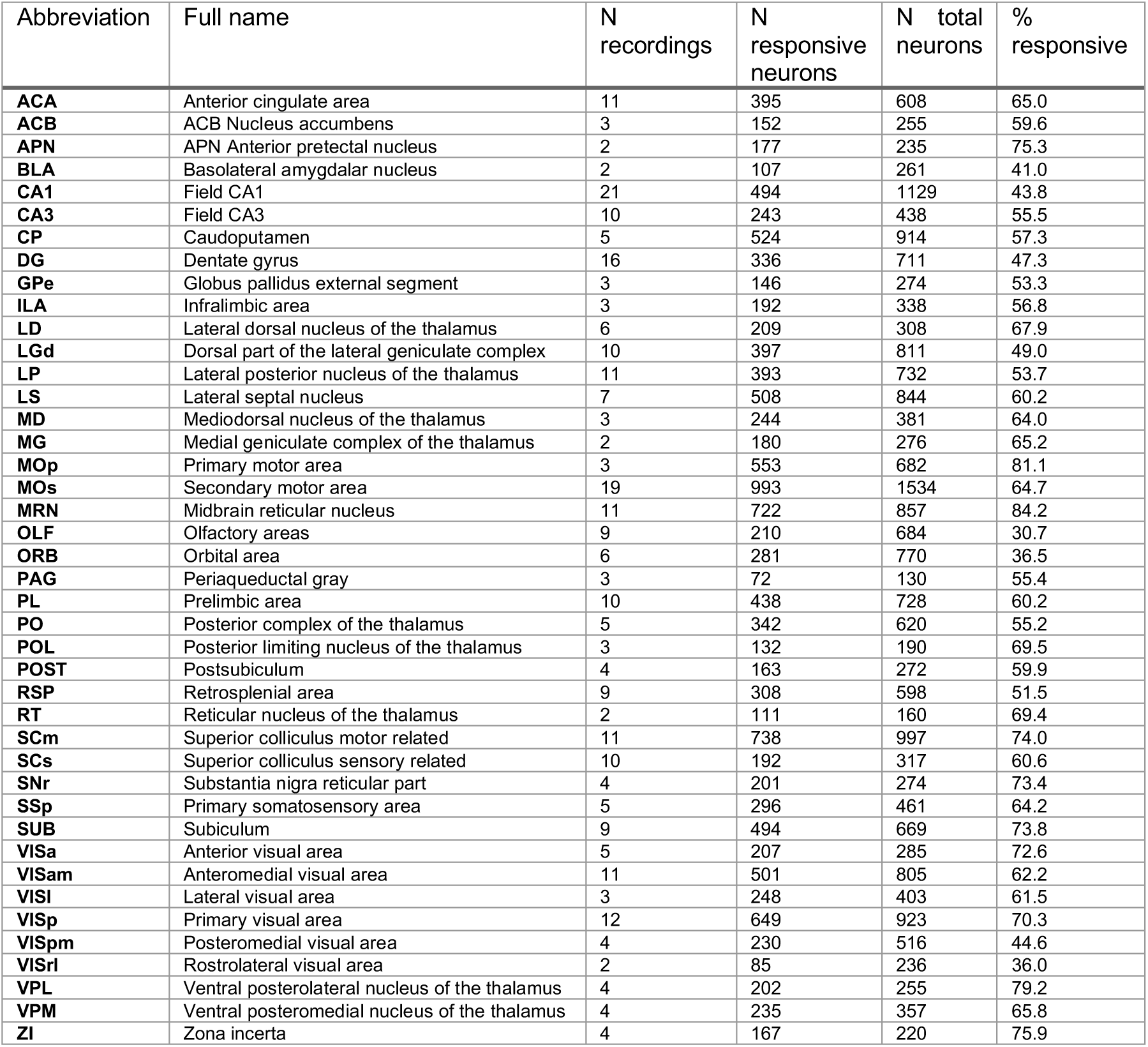
Brain regions recorded

**Extended Data Table 2.** Subjects included in the study

| **Subject ID** | **Sex** | **Genotype** |
| --- | --- | --- |
| **Cori** | F | tetO-G6s x CaMK-tTA |
| **Forssmann** | M | C57BL/6J |
| **Hench** | M | tetO-G6s x CaMK-tTA |
| **Lederberg** | F | Vglut1-IRES2-Cre-D |
| **Moniz** | M | RCL-GCaMP6f x VGlut-Cre |
| **Muller** | F | RCL-GCaMP6f x VGlut-Cre |
| **Radnitz** | F | Snap25-GCaMP6s |
| **Richards** | M | C57BL/6J |
| **Tatum** | F | Vglut1-IRES2-Cre-D |
| **Theiler** | F | RCL-GCaMP6f x VGlut-Cre |

**Extended Data Table 3.**
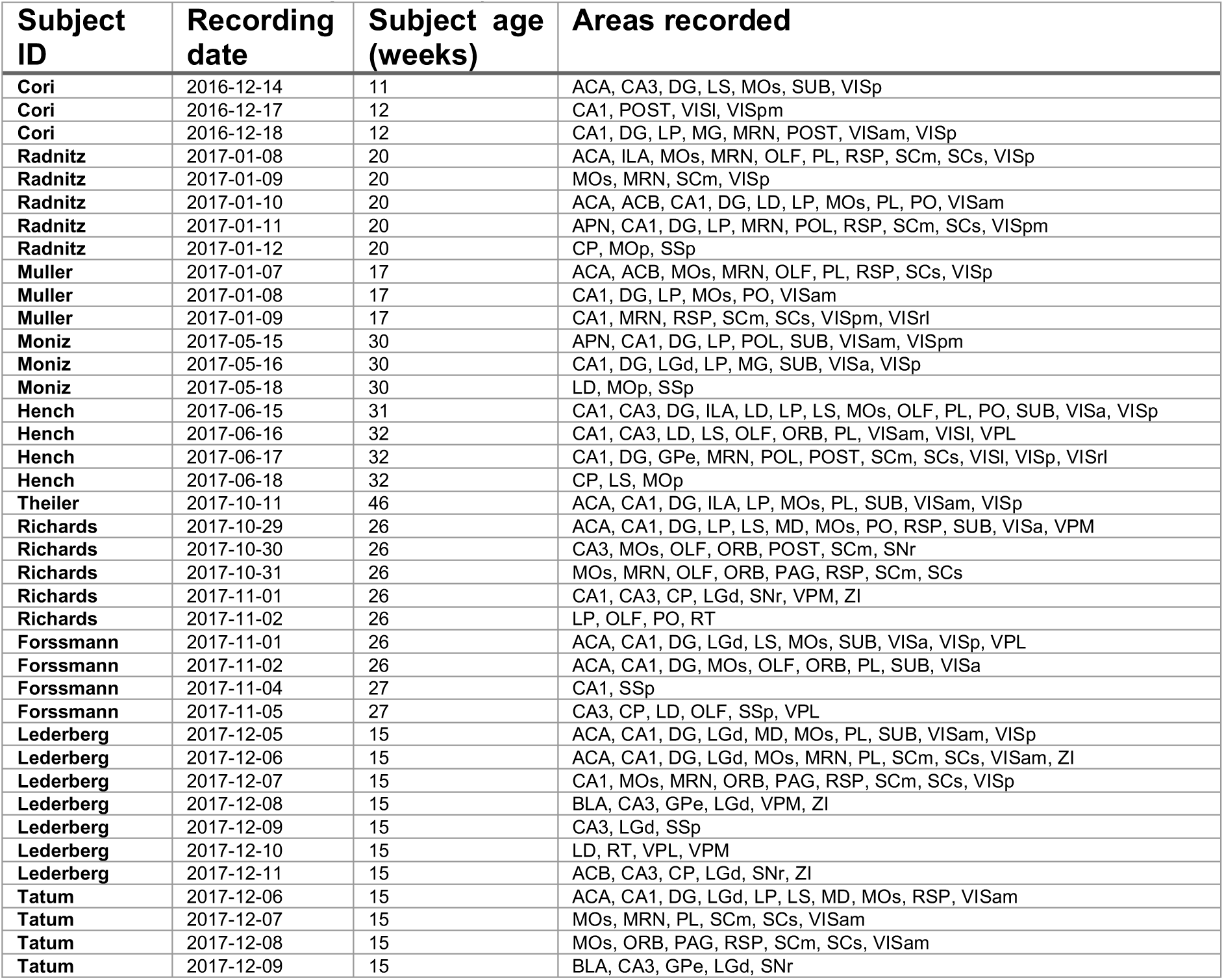
Recording sessions

| **Subject ID** | **Recording date** | **Subject age (weeks)** | **Areas recorded** |
| --- | --- | --- | --- |
| Cori | 2016-12-14 | 11 | ACA, CA3, DG, LS, MOs, SUB, VISp |
| Cori | 2016-12-17 | 12 | CA1, POST, VISl, VISpm |
| Cori | 2016-12-18 | 12 | CA1, DG, LP, MG, MRN, POST, VISam, VISp |
| Radnitz | 2017-01-08 | 20 | ACA, ILA, MOs, MRN, OLF, PL, RSP, SCm, SCs, VISp |
| Radnitz | 2017-01-09 | 20 | MOs, MRN, SCm, VISp |
| Radnitz | 2017-01-10 | 20 | ACA, ACB, CA1, DG, LD, LP, MOs, PL, PO, VISam |
| Radnitz | 2017-01-11 | 20 | APN, CA1, DG, LP, MRN, POL, RSP, SCm, SCs, VISpm |
| Radnitz | 2017-01-12 | 20 | CP, MOp, SSp |
| Muller | 2017-01-07 | 17 | ACA, ACB, MOs, MRN, OLF, PL, RSP, SCs, VISp |
| Muller | 2017-01-08 | 17 | CA1, DG, LP, MOs, PO, VISam |
| Muller | 2017-01-09 | 17 | CA1, MRN, RSP, SCm, SCs, VISpm, VISrl |
| Moniz | 2017-05-15 | 30 | APN, CA1, DG, LP, POL, SUB, VISam, VISpm |
| Moniz | 2017-05-16 | 30 | CA1, DG, LGd, LP, MG, SUB, VISa, VISp |
| Moniz | 2017-05-18 | 30 | LD, MOp, SSp |
| Hench | 2017-06-15 | 31 | CA1, CA3, DG, ILA, LD, LP, LS, MOs, OLF, PL, PO, SUB, VISa, VISp |
| Hench | 2017-06-16 | 32 | CA1, CA3, LD, LS, OLF, ORB, PL, VISam, VISl, VPL |
| Hench | 2017-06-17 | 32 | CA1, DG, GPe, MRN, POL, POST, SCm, SCs, VISl, VISp, VISrl |
| Hench | 2017-06-18 | 32 | CP, LS, MOp |
| Theiler | 2017-10-11 | 46 | ACA, CA1, DG, ILA, LP, MOs, PL, SUB, VISam, VISp |
| Richards | 2017-10-29 | 26 | ACA, CA1, DG, LP, LS, MD, MOs, PO, RSP, SUB, VISa, VPM |
| Richards | 2017-10-30 | 26 | CA3, MOs, OLF, ORB, POST, SCm, SNr |
| Richards | 2017-10-31 | 26 | MOs, MRN, OLF, ORB, PAG, RSP, SCm, SCs |
| Richards | 2017-11-01 | 26 | CA1, CA3, CP, LGd, SNr, VPM, ZI |
| Richards | 2017-11-02 | 26 | LP, OLF, PO, RT |
| Forssmann | 2017-11-01 | 26 | ACA, CA1, DG, LGd, LS, MOs, SUB, VISa, VISp, VPL |
| Forssmann | 2017-11-02 | 26 | ACA, CA1, DG, MOs, OLF, ORB, PL, SUB, VISa |
| Forssmann | 2017-11-04 | 27 | CA1, SSp |
| Forssmann | 2017-11-05 | 27 | CA3, CP, LD, OLF, SSp, VPL |
| Lederberg | 2017-12-05 | 15 | ACA, CA1, DG, LGd, MD, MOs, PL, SUB, VISam, VISp |
| Lederberg | 2017-12-06 | 15 | ACA, CA1, DG, LGd, MOs, MRN, PL, SCm, SCs, VISam, ZI |
| Lederberg | 2017-12-07 | 15 | CA1, MOs, MRN, ORB, PAG, RSP, SCm, SCs, VISp |
| Lederberg | 2017-12-08 | 15 | BLA, CA3, GPe, LGd, VPM, ZI |
| Lederberg | 2017-12-09 | 15 | CA3, LGd, SSp |
| Lederberg | 2017-12-10 | 15 | LD, RT, VPL, VPM |
| Lederberg | 2017-12-11 | 15 | ACB, CA3, CP, LGd, SNr, ZI |
| Tatum | 2017-12-06 | 15 | ACA, CA1, DG, LGd, LP, LS, MD, MOs, RSP, VISam |
| Tatum | 2017-12-07 | 15 | MOs, MRN, PL, SCm, SCs, VISam |
| Tatum | 2017-12-08 | 15 | MOs, ORB, PAG, RSP, SCm, SCs, VISam |
| Tatum | 2017-12-09 | 15 | BLA, CA3, GPe, LGd, SNr |

### Surgery

A brief (~1 h) initial surgery was performed under isoflurane (1-3% in O2) anesthesia to implant a steel headplate (~15 x 3 x 0.5mm, ~1 g) and, in most cases, a 3D-printed recording chamber. The chamber was a semi-conical, opaque piece of polylactic acid (PLA) with 12 mm diameter upper surface, and lower surface designed to fit to the shape of an average mouse skull, exposing approximately the area from 3.5 anterior to 5.5 posterior to bregma, and 4.5 left to 4.5 right, and narrowing near the eyes. The implantation method largely followed the method of Guo et al^46^ with some modifications and was described previously^45^. In brief, the dorsal surface of the skull was cleared of skin and periosteum and prepared with a brief application of green activator (Super-Bond C&B, Sun Medical Co.). The chamber was attached to the skull with cyanoacrylate (VetBond; World Precision Instruments) and the gaps between the cone and the skull were filled with L-type radiopaque polymer (Super-Bond C&B). A thin layer of cyanoacrylate was applied to the skull inside the cone and allowed to dry. Thin layers of UV-curing optical glue (Norland Optical Adhesives #81, Norland Products) were applied inside the cone and cured until the exposed skull was covered. The headplate was attached to the skull over the interparietal bone with Super-Bond polymer, and more polymer was applied around the headplate and cone.

Following recovery, mice were given three days to recover while being treated with carprofen, then acclimated to handling and head-fixation prior to training.

### Two-alternative unforced choice task

The two-alternative unforced choice task design was described previously^31^. In this task, mice were seated on a plastic apparatus with forepaws on a rotating wheel, and were surrounded by three computer screens (Adafruit, LP097QX1) at right angles covering 270 x 70 degrees of visual angle (d.v.a.). Each screen was ~11cm from the mouse’s eyes at its nearest point and refreshed at 60Hz. The screens were fitted with Fresnel lenses (Wuxi Bohai Optics, BHPA220-2-5) to ameliorate reductions in luminance and contrast at larger viewing angles near their edges, and these lenses were coated with scattering window film (“frostbite”, The Window Film Company) to reduce reflections. The wheel was a ridged rubber Lego wheel affixed to a rotary encoder (Kubler 05.2400.1122.0360). A plastic tube for delivery of water rewards was placed near the subject’s mouth. Licking behavior was monitored by attaching a piezo film (TE Connectivity, CAT-PFS0004) to the plastic tube and recording its voltage. Eye movements were monitored by illuminating the subject with IR light (830nm, Mightex SLS-0208-A) and monitoring the right eye with a camera (The Imaging Source, DMK 23U618) fitted with zoom lens (Thorlabs MVL7000) and long-pass filter (Thorlabs FEL0750). Body movements were monitored with another camera situated above the central screen. Full details of the experimental apparatus including detailed parts list can be found at http://www.ucl.ac.uk/cortexlab/tools/wheel.

A trial was initiated after the subject had held the wheel still for a short interval (duration uniformly distributed between 0.2-0.5 sec on each trial; Figure 1b). At trial initiation, a visual stimulus was presented on the left, right, both, or neither screen. The stimulus was a gabor patch with orientation 45 degrees, sigma 9 d.v.a., and spatial frequency 0.1 cycles/degree. After stimulus onset there was a random delay interval of 0.5-1.2 sec, during which time the subject could turn the wheel without penalty, but visual stimuli were locked in place and rewards could not be earned. The subjects nevertheless typically responded immediately to the stimulus onset. At the end of the delay interval, an auditory go cue was delivered (8 kHz pure tone for 0.2 sec) after which the visual stimulus position became coupled to movements of the wheel. Wheel turns in which the top surface of the wheel was moved to the subject’s right led to rightward movements of stimuli on the screen, i.e. a stimulus on the subject’s left moved towards the central screen. Put another way, clockwise turns of the wheel, from the perspective of the mouse, led to clockwise movement of the stimuli around the subject. A left or right turn was registered when the wheel was turned by an amount sufficient to move the visual stimuli by 90 d.v.a. in either direction. When at least one stimulus was presented, the subject was rewarded for driving the higher contrast visual stimulus to the central screen (if both stimuli had equal contrast, left/right turns were rewarded with 50% probability). When no stimuli were presented, the subject was rewarded if no turn was registered during the 1.5 s following the go cue. Immediately following registration of a choice or expiry of the 1.5 s window, feedback was delivered. If correct, feedback was a water reward (2 – 3 µL) delivered by the opening of a valve on the water tube for a calibrated duration. If incorrect, feedback was a white noise sound played for 1 s. During the 1 s feedback period, the visual stimulus remained on the screen. After a subsequent inter-trial interval of 1 s, the mouse could initiate another trial by again holding the wheel still for the prescribed duration.

Mice were trained on this task with the following shaping protocol. First, high contrast stimuli (50 or 100%) were presented only on the left or the right, with an unlimited choice window, and repeating trial conditions following incorrect choices (‘repeat on incorrect’). Once mice achieved high accuracy and initiated movements rapidly – approximately 70 or 80% performance on non-repeat trials, and with reaction times nearly all < 1 second, but at the experimenter’s discretion – trials with no stimuli were introduced, again repeating on incorrect. Once subjects responded accurately on these trials (70 or 80% performance, at experimenter’s discretion), lower contrast trials were introduced without repeat on incorrect. Finally, contrast comparison trials were introduced, starting with high vs low contrast, then high vs medium and medium vs low, then trials with equal contrast on both sides. The final proportion of trials presented was weighted towards easy trials (high contrast vs zero, high vs low, medium vs zero, and no-stimulus trials) to encourage high overall reward rates and sustained motivation.

On most trials for which subjects eventually made a left or right turn by the end of the trial, the subjects responded immediately to the stimulus presentation, turning the wheel within 400 ms of stimulus appearance (64.9 ± 14.0% s.d., n=39 sessions), nearly always in the same direction as their final choice (96.6 ± 3.4%). For this study, data analyses focused on this initial 400 ms period, and we defined Left and Right Choice trials as in which this period contained a clockwise or counterclockwise turn of sufficient amplitude (90 d.v.a.), and NoGo trials as those where it contained no detectable movement. To exclude trials in which wheel turns were coincidentally made before subjects could respond to the stimuli, only trials with movement onset between 125 to 400ms post-stimulus onset, or with no movement of any kind during the window from −50 to 400ms post-stimulus onset, were included. Trials with other movements, that were detectable but would not have resulted in registering a choice by the end of the movement, were excluded.

Behavioral trials when the mouse was disengaged were excluded from analysis. These trials were defined as Miss trials (stimulus present but wheel not turned) preceded by two or more other Miss trials, as well as all NoGo trials occurring consecutively at the end of the session.

When analyzing activity following reward delivery, only correct NoGo trials were included, i.e. trials with no visual stimulus and no wheel movement.

Sessions were included when at least 12 trials of each type (Left, Right, NoGo) could be included for analysis, and when anatomical localization was sufficiently confident (see below).

For analyses requiring trials with different choices but matched for stimulus contrast, we considered all trials with contralateral stimulus contrast greater than zero, and split them by low, medium, and high contralateral contrast. For each contrast level, we counted the number of trials with that contrast and each response type (Left or Right; NoGo; or passive condition). We took the minimum of these three numbers, and selected that many trials randomly from each group. This resulted in three sets of trials-trials with Left or Right choices; trials with NoGos; and trials in the passive condition – which each contained low, medium, and high contralateral contrasts but which all contained exactly the same numbers of each contrast. When fewer than 10 such trials could be found, the session was excluded for the matched-contrast analyses (n=34 of 39 sessions included).

### Neuronal recordings

Recordings were made using Neuropixels (“Phase3A”) electrode arrays^30^, which have 384 selectable recording sites out of 960 sites on a 1 cm shank. Probes were mounted to a custom 3D-printed PLA piece and affixed to a steel rod held by a micromanipulator (uMP-4, Sensapex Inc.). To allow later track localization, prior to insertion probes were coated with a solution of DiI (ThermoFisher Vybrant V22888 or V22885) by holding 2µL in a droplet on the end of a micropipette and touching the droplet to the probe shank, letting it dry, and repeating until the droplet was gone, after which the probe appeared pink.

On the day of recording or within two days before, mice were briefly anaesthetized with isoflurane while one or more craniotomies were made, either with a dental drill or a biopsy punch. After at least three hours of recovery, mice were head-fixed in the setup. Probes had a soldered connection to short external reference to ground; the ground connection at the headstage was subsequently connected to an Ag/AgCl wire positioned on the skull. The craniotomies as well as the wire were covered with saline-based agar. The agar was covered with silicone oil to prevent drying. In some experiments a saline bath was used rather than agar. Probes were advanced through the agar and through the dura, then lowered to their final position at ~10µm/sec. Electrodes were allowed to settle for ~15 min before starting recording. Recordings were made in external reference mode with LFP gain = 250 and AP gain = 500. Recordings were repeated at different locations on each of multiple subsequent days, performing new craniotomy procedures as necessary. All recordings were made in the left hemisphere.

### Passive stimulus presentation

After each behavior session we performed a passive replay experiment while continuing to record from the same electrodes. Mice were presented with two types of sensory stimuli without possibility of receiving reward for any behavior: replay of task stimuli; and sparse flashed visual stimuli for receptive field mapping.

The replayed task stimuli were: left and right visual stimuli of each contrast; some combinations of left and right visual stimuli simultaneously; go cue beeps; white noise bursts; and reward valve clicks (but with a manual valve closed so that no water was delivered). These stimuli were replayed at 1-2 sec randomized intervals for 10 or 25 randomly interleaved repetitions each.

Receptive fields were mapped with white squares of 8 d.v.a. edge length, positioned on a 10 x 36 grid (some stimulus positions were located partially off-screen) on a black background. The stimuli were shown for 10 monitor frames (167ms) at a time, and their times of appearance were independently randomly selected to yield an average rate of ~0.12 Hz.

### Data analysis

The data were automatically spike sorted with Kilosort^33^ (https://github.com/cortex-lab/Kilosort) and then manually curated with the ‘phy’ gui (https://github.com/kwikteam/phy). Extracellular voltage traces were preprocessed common-average referencing^47^: subtracting each channel’s median to remove baseline offsets, then subtracting the median across all channels at each time point to remove artifacts. During manual curation, each set of events (‘unit’) detected by a particular template was inspected and if the events (‘spikes’) comprising the unit were judged to correspond to noise (zero or near-zero amplitude; non-physiological waveform shape or pattern of activity across channels), the entire unit was discarded. Units containing low-amplitude spikes, spikes with inconsistent waveform shapes, and/or refractory period contamination were labeled as ‘multi-unit activity’ and not included for further analysis. Finally, each unit was compared to similar, spatially neighboring units to determine whether they should be merged, based on spike waveform similarity, drift patterns, or cross-correlogram features. Units were also excluded if their average rate in the analysis window (stimulus onset to 0.4 sec after; ‘trial firing rate’) was less than 0.1 Hz. Units passing these criteria were considered to reflect the spiking activity of a neuron.

Neurons were only included for further analysis least 13 neurons passing the above criteria were identified as coming from the same brain region, in the same experiment. Furthermore, brain regions were only included for which at least two recordings had sufficient numbers of neurons.

To determine whether a neuron was active during the task (Fig. 1i), a set of six statistical tests were used to detect changes in activity during various task epochs and conditions: 1) Wilcoxon signrank test between trial firing rate (rate of spikes between stimulus onset and 400 ms post-stimulus) and baseline rate (defined in period −0.2 to 0 s relative to stimulus onset on each trial); 2) signrank test between stimulus driven rate (firing rate between0.05 and 0.15 s after stimulus onset) and baseline rate; 3) signrank test between pre-movement rates (−0.1 to 0.05 s relative to movement onset) and baseline rate (for trials with movements); 4) Wilcoxon ranksum test between pre-movement rates on left choice trials and those on right choice trials; 5) signrank test between post-movement rates (−0.05 to 0.2 s relative to movement onset) and baseline rate; 6) ranksum test between post-reward rates (0 to 0.15 s relative to reward delivery for correct NoGos) and baseline rates. A neuron was considered active during the task, or to have detectable modulation during some part of the task, if any of the p-values on these tests were below a Bonferroni-corrected alpha value (0.05/6 = 0.0083). However, because the tests were coarse and would be relatively insensitive to neurons with transient activity, a looser threshold was used to determine the neurons included for statistical analyses (Figs. 3-5): if any of the first four tests (i.e. those concerning the period between stimulus onset and movement onset) had a p-value less than 0.05.

For visualizing PSTHs (Figs. 2 and 3c), the activity of each neuron was then binned at 0.005 s, smoothed with a causal half-Gaussian filter with standard deviation 0.02 s, averaged across trials, smoothed with another causal half-gaussian filter with standard deviation 0.03 s, baseline subtracted (baseline period −0.02 to 0 s relative to stimulus onset, including all trials in the task), and divided by baseline + 0.5 sp/s. Neurons were selected for PSTH display if they had a significant difference between firing rates on trials in the task with stimuli and movements versus those without both, using a sliding window 0.1 s wide and in steps of 0.005 s (ranksum p<0.0001 for at least three consecutive bins).

Visual receptive fields were determined by sparse noise mapping outside the context of the behavioral task. The evoked rates for each presentation were measured as the spike count in the 200ms following stimulus onset. The rates evoked by stimuli at the peak location and surrounding four nearest locations were combined and compared to the rates for all locations > 45 d.v.a. from the peak location using a Wilcoxon ranksum test. Any neurons for which the p value of the test was less than 10^-6^ were counted as having a significant visual receptive field. If the peak receptive field location was within 18 d.v.a. of the location used for the visual stimulus in the behavioral task (i.e. within 2x the standard deviation of the Gaussian aperture of that stimulus), the neuron was counted as having an ‘on-target’ receptive field.

To statistically compare pre-trial firing rates between the active and passive conditions (i.e. between trials of active task performance, versus later passive stimulus replay, Fig. 3b), we performed a nested multiple ANOVA test, in order to account for correlated variability between neurons within recording sessions. Each observation was a neuron’s average measured pre-trial firing rate in the window between 250 and 50ms prior to stimulus onset, log transformed (log_10_(*x* + 1 sp/s)) to make distributions approximately normal. Any trials with detectable wheel movement in this interval were excluded. The ANOVA had three factors: active/passive condition, recording session, and neuron identity (nested within recording session). The null hypothesis of no difference between baseline rates in active and passive conditions for neurons from a given brain region was rejected if the p-value for the active/passive condition factor was less than 0.0012, i.e. less than 0.05 after applying a Bonferroni correction for the 42 brain regions tested.

To compute the trial-by-trial ‘engagement index’, we took the difference in pre-stimulus firing between the average of all task (‘active’) trials a and of all passive trials p,

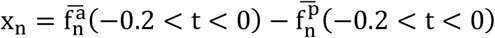

This quantity was computed for each neuron in one of the areas with significant differences between task and passive determined by nested ANOVA analysis (319 ± 32.5 mean ± s.e. neurons included per session, n=34 sessions), and accumulated into a vector *x* for each session. We normalized each session’s vector to unit magnitude. We computed the dot product **x** ⋅ **f**^**i**^ of *x* with the vector of pre-stimulus firing rates for each trial *i*. We then computed the mean across movement and the mean across miss trials, and took the difference of the two.

### Reduced rank kernel regression

To identify choice-selective neurons, we began by fitting a ‘kernel regression’ model^34,48,49^. In this analysis, the firing rate of each neuron is described as a linear sum of temporal filters aligned to task events. For the current study, only visual stimulus onset and wheel movement onset kernels were required, since we consider here only the period in between the two. In the model, the predicted firing rate 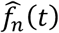 for neuron *n* is given as

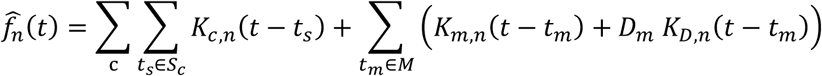

Here, *c* represents of the the 6 stimulus types (contralateral low, medium, or high, or ipsilateral low, medium, or high), *S*_*c*_ represents the set of times for which this contrast appeared, and *K*_*c,n*_(*t*) represents the Stimulus kernel function of this contrast for neuron *n*. *M* represents the set of movement times and *K*_*m,n*_(*t*) represents the Movement kernel for neuron *n*; *D*_m_ represents direction of movement *m* (encoded as ±1), and *K*_T,B_ represents the Choice kernel for neuron *n*. The Stimulus kernels *K*_*c,n*_(*t*) are supported over the window −0.05 to 0.4 s relative to stimulus onset, and the Movement and Choice kernels are supported over the window 0.25 to 0.025 s relative to movement onset. Prior to estimating the kernels, the discretized firing rates *f*_*n*_(*t*) for each neuron were estimated by binning spikes into 0.005 s bins and then smoothing with a causal half-Gaussian filter with standard deviation 0.025 s. The stimulus kernels therefore contain *L*_*c*_ = 90 time bins, while movement kernels contain *L*_*d*_ = 55 time bins.

The large number of parameters to be fit, combined with the relatively small number of trials of each type pose a challenge for estimation. We devised a solution to this problem that leverages the large number of neurons recorded using reduced rank regression.

First, for each kernel to be fit, we construct a Toeplitz predictor matrix. For stimuli of contrast *c*, we define a Toeplitz predictor matrix *P*_*c*_ of size *T* × *L*_*c*_, where *T* is the total number of time points in the training set, and *L*_*c*_ is the number of lags required for the stimulus kernels. The predictor matrix contains diagonal stripes starting each time a stimulus of contrast *c* is presented: *P*_*c*_(*t,l*) = 1 if *t* − *l* ∈ *S*_*c*_ and 0 otherwise. Predictor matrices for the Movement and Choice kernels are defined similarly, and the six stimulus predictor matrices and two movement predictors are horizontally concatenated to yield a global prediction matrix *P* of size *T*× 650 ⋅ (650 = 6*L*_*c*_ + 2*L*_*d*_ is the total length of all kernels for one neuron.)

The simplest approach to fit the kernel shapes would be to minimize the squared error between true and predicted firing rate using linear regression. To do this, we would horizontally concatenating the rate vectors of all *N* neurons together into a *T* × *N* matrix *F*, and estimate the kernels for each neuron by finding a matrix *K* of size 650 × *N* to minimize the squared error:

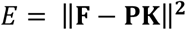

However, as each *k*_*n*_ has 650 parameters, linear regression results in noisy and overfit kernels when fit to a single neuron, particularly given the high trial-to-trial variability of neuronal firing. Although expressing the kernels as a sum of basis functions can reduce the number of required parameters^34^, the success of this method depends strongly on the choice of basis functions, with an appropriate choice will differ depending on properties of the task and stimuli. However, as each neuron’s kernel has 650 parameters, linear regression results in noisy and overfit kernels when fit to a single neuron, particularly given the high trial-to-trial variability of neuronal firing. Although expressing the kernels as a sum of basis functions can reduce the number of required parameters^34^, the success of this method depends strongly on the choice of basis functions, with an appropriate choice will differ depending on properties of the task and stimuli. The large number of neurons in the current dataset allows an alternative approach.

This approach is based on reduced rank regression^50^, which allows regularized estimation by factorizing the kernel matrix **K** into the product of a 650 × 650 matrix **B** and 650 × *N* matrix **W** minimizing the total error:

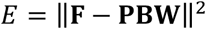

The *T* × *r* matrix *PB* may be considered as a set of temporal basis functions, which can be linearly combined to estimate each neuron’s firing rate over the whole training set. Reduced rank regression ensures that these basis functions are ordered, so that predicting population activity from only the first *r* columns will result in the best possible prediction from any rank *r* matrix.

To estimate each neuron’s kernel functions, we estimated a weight vector *w*_*n*_ to minimize an error *E*_*n*_ = |*f*_*n*_ − *PBw*_*n*_|^2^ for each neuron with elastic net regularization (using the package cvglmnet for Matlab with parameters α = 0.5 and λ = 0.5), and used cross-validation to determine the optimal number of columns *r*_*n*_ of *PB* to keep when predicting neuron *n*. The kernel functions for neuron *n* were then unpacked from the 650-dimensional vector obtained by multiplying the first *r*_*n*_ columns of *B* by *w*_*n*_. Neurons with maximal cross-validated variance explained of <2% were excluded from subsequent analyses.

To assess the selectivity of individual neurons for each kernel, we first fit the activity of each neuron again using the reduced rank regression procedure above (including deriving a new basis set) but excluding the kernel to be tested. We subtracted this prediction from the raw data to yield residuals, representing aspects of the neuron’s activity not explainable from the other kernels. We then repeated the reduced rank regression procedure one more time, using the residual firing rates as the independent variable, and using only the test kernel. The cross-validated quality of this fit determined the variance explainable only by the test kernel. If this variance explained was >2%, the neuron was deemed selective for that kernel and was included in Figure 4b-d.

To perform population decoding, we began with the residual firing rates produced as described above, produced by fitting without a test kernel. We then split trials in a binary fashion: trials that had vs. did not have an ipsilateral stimulus; had vs. did not have a contralateral stimulus; had vs. did not have any movement (either left or right); had left choice vs. had right choice (considering only trials with one of the two). We identified a population coding direction encoding the difference between the two sets of trials, by fitting an L1-regularized logistic regression on data from training trials, using the period 0.05 to 0.15 sec relative to stimulus onset for Stimulus decoding, and the period −0.05 to 0 sec relative to movement onset for Move or Choice decoding. We then predicted the binary category of test data by projecting firing rates from test set trials, from each time point during the trial, onto the weight vector of the logistic regression. The population decoding was taken as the difference between projections between test set trials of each binary category.

To compare decoding timecourse across areas, we took the population decoding from each key area in each recording (n=29 populations from frontal cortex including MOs, PL, and MOp; n=29 populations from midbrain including SCm, MRN, SNr, ZI; n = 5 striatum, CP), normalized each so that the mean across recordings within an area was 1, and performed a two-way ANOVA. The two factors were time relative to movement onset and area (frontal, midbrain, striatum). We found a significant effect of time (50 d.f., F = 10.43, p<10^-71^) but no significant effect of area (2 d.f., F = 0.28, p>0.05) and no significant interaction between time and area (100 d.f., F = 0.12, p>0.05).

### Anatomical targeting and probe localization

To determine probe insertion trajectories, we first selected desired recording sites, and then designed appropriate trajectories using the allen_atlas_probe gui (A. J. Peters, www.github.com/cortex-lab/allenCCF). In doing so, Allen CCF coordinate [5.4 mm AP, 0 DV, 5.7 LR] was taken as the location of bregma. Craniotomies were targeted accordingly and angles of insertion were set manually. For some of the visual cortex recordings, surface insertion coordinates were targeted based on prior widefield calcium imaging. Using techniques described previously^45^, we imaged activity across cortex during presentation of the sparse visual noise receptive field mapping stimulus described above. Responses to visual stimuli near the intended location of the task stimuli were combined and used to identify cortical locations with retinotopically-aligned neurons. In some cases, these same imaging sessions were used to target MOp and SSp recordings to the area of large activity observed during forelimb movements that covers both of those areas^51^. Finally, MOs recordings were targeted at and around the cortical coordinates identified as disrupting task performance when inactivated, around +2 mm AP, 1 mm ML^51^.

Recording sites were localized to brain regions by manual inspection of histologically identified recording tracks, in combination with alignment to the Allen Institute Common Coordinate Framework, as follows.

Mice were perfused with 4% PFA, the brain was extracted and fixed for 24 hours at 4 C in PFA, then transferred to 30% sucrose in PBS at 4 C. The brain was mounted on a microtome in dry ice and sectioned at 60 µm slice thickness. Sections were washed in PBS, mounted on glass adhesion slides, and stained with DAPI (Vector Laboratories, H-1500). Images were taken at 4x magnification for each section using a Zeiss AxioScan, in three colors: blue for DAPI, green for GCaMP (when present), and red for DiI.

An individual DiI track was typically visible across multiple slices, and recording locations along the track were manually identified by comparing structural aspects of the histological slice with features in the atlas. In most cases, this identification was aided by reconstruction of the track in Allen CCF coordinates. To achieve this, a manual initial guess was made of the 3D Allen CCF coordinate for each observed DiI mark. In some cases, this guess was aided by a control-point registration of the histological slice to an atlas slice. Once the 3D coordinates were identified for each DiI mark along the track, a line was fitted to these coordinates in 3D and the atlas labels were extracted from along this line. Together, these approaches resulted in identification of the list of brain regions each probe track passed through.

We observed idiosyncratic scaling of each brain relative to the size of the atlas, and also that the electrode tip was difficult to precisely localize in the histological data. To overcome these problems, in a final step, the set of brain regions along the probe track was aligned to the recording sites by use of physiological signatures. For instance, segments of the probe that were located in ventricles, fiber tracts, or dendrite layers of the hippocampus were readily identifiable due to the low observed spike rate. Other identifiers included LFP signatures of hippocampus and olfactory areas, firing rates and spike amplitudes for cortical layers and certain subcortical areas, and presence of features such as visual receptive fields for superficial layers of SC and for visual thalamic nuclei. Such signatures were used to find a scaling and shift that aligned the list of identified brain regions along the track to recorded positions on the probe. Data recorded during the behavioral task was not considered during this alignment procedure.

